# Structural insights into ligand recognition and selectivity of the human hydroxycarboxylic acid receptor HCAR2

**DOI:** 10.1101/2023.03.28.534513

**Authors:** Xin Pan, Fang Ye, Peiruo Ning, Zhiyi Zhang, Binghao Zhang, Geng Chen, Wei Gao, Chen Qiu, Zhangsong Wu, Kaizheng Gong, Jiancheng Li, Jiang Xia, Yang Du

## Abstract

Hydroxycarboxylic acid receptor 2 (HCAR2) belongs to the family of class A G-protein-coupled receptors with key roles in regulating lipolysis and free fatty acid formation in humans. It is deeply involved in many pathophysiological processes and serves as an attractive target for the treatment of neoplastic, autoimmune, neurodegenerative, inflammatory, and metabolic diseases. Here, we report four cryo-EM structures of human HCAR2-Gi1 complexes with or without agonists, including the drugs niacin and acipimox, and the highly subtype-specific agonist MK-6892. Combined with molecular docking and functional analysis, we have revealed the recognition mechanism of HCAR2 for different agonists and summarized the general pharmacophore features of HCAR2 agonists, which are based on three key residues R111^3.36^, S179^45.52^, and Y284^7.43^. Notably, the MK-6892-HCAR2 structure shows an extended binding pocket relative to other agonist-bound HCAR2 complexes. In addition, the key residues that determine the ligand selectivity between the HCAR2 and HCAR3 are also illuminated. Our findings provide structural insights into the ligand recognition, selectivity, activation, and G protein coupling mechanism of HCAR2, which sheds light on the design of new HCAR2-targeting drugs for greater efficacy, higher selectivity, and fewer or no side effects.

## 1 Introduction

Hydroxy carboxylic acid receptor 2 (HCAR2), also known as GPR109A, is an important metabolite-sensing receptor present in most mammalian species, and belongs to the class A G-protein-coupled receptor (GPCR) family.^1–3^ HCAR2 is highly expressed in multiple cell types (e.g., adipocytes, immune cells, retinal pigmented cells, colonic epithelial cells, keratinocytes, and microglia) and mediates downstream signaling transducers by coupling to the Gi/o family of G-proteins.^4, 5^ The endogenous ligands of HCAR2 are β-hydroxybutyrate (β-OHB) and butyrate, both of which serve as nutrient sources for cells under various physiological conditions.^2, 6^ In particular, upon starvation or other extreme conditions, HCAR2 is activated by elevated β-OHB in vivo to reduce the lipolysis and free fatty acid formation in adipocytes, thus promoting efficient utilization of fat energy stores and preventing the development of ketoacidosis.^7, 8^ Moreover, HCAR2 is implicated in mitigating many pathophysiological processes, including the reduction of chemokine and proinflammatory cytokine production; amelioration of atherosclerosis, sepsis, and diabetic retinopathy; suppression of the occurrence of breast cancer, colitis, and acute pancreatitis; and maintenance of the integrity of the intestinal barrier.^5, 9, 10^ Emerging studies also indicate that the activation of microglial HCAR2 has beneficial effects on multiple neurological disorders, such as Alzheimer’s disease, Parkinson’s disease, multiple sclerosis, stroke, and pathological pain conditions.^11, 12^ Collectively, HCAR2 is emerging as an attractive therapeutic target for the treatment of neoplastic, autoimmune, neurodegenerative, inflammatory, and metabolic diseases.

Currently, several highly potent HCAR2 agonists, including niacin, acipimox, acifran, and monomethyl fumarate (MMF), have been approved for clinical treatment of cardiovascular and neurological disorders.^13^ Of these, niacin, serves as a well-known agonist of HCAR2. It is the oldest lipid-lowering drug to date, resulting from its ability to lower very low-density lipoprotein cholesterol, low-density lipoprotein cholesterol, and lipoprotein levels, as well as to increase high-density lipoprotein cholesterol levels to a greater extent than many other marketed drugs.^14^ In addition, niacin has well-established antiatherogenic effects which, to a considerable degree, are based on the HCAR2-mediated anti-inflammatory effects of reducing the M1 macrophage proportion.^15, 16^ The latest research also shows that niacin is being investigated as a drug for Parkinson’s disease and glioblastoma because of its immunomodulatory and neuroprotective properties, and clinical trials are currently in progress (NCT03808961 and NCT04677049).^17^ The niacin-derived antilipolytic drugs acipimox and acifran are generally used to treat dyslipidemia and atherosclerosis clinically.^18^ Moreover, MMF was approved by the FDA in 2020 for the treatment of relapsing multiple sclerosis.^11, 19^ It has been demonstrated that HCAR2 at least in part mediates the beneficial effects of MMF. This was confirmed by a study of Parodi et al., in which MMF switched the LPS-activated microglia from a pro-inflammatory type to a neuroprotective one through activation of HCAR2.^20^ Overall, the agents targeting HCAR2 have achieved notable successes in treating a variety of clinical diseases; nevertheless, several important challenges still remain. First, despite the good treatment efficacy of niacin, acipimox, and acifran, their use is less widespread than statins for the treatment lipid disorders, which is mainly attributed to an uncomfortable cutaneous flushing effect that limits patient compliance.^21^ Given this, some highly subtype-specific HCAR2 agonists (e.g., MK-6892, SCH900271, and GSK256073) have been developed, which share the lipid-lowering effects, but significantly alleviate the flushing effect.^22–24^ This leads us to question as to what the structural differences are between these subtype-specific agonists and approved drugs when bound to HCAR2? So far, no experimental structures have yet been resolved for any hydroxycarboxylic acid receptor family, including HCAR1–3. Therefore, the detailed binding modes and recognition mechanisms of endogenous ligands, therapeutic agents, and certain subtype-specific HCAR2 agonists are largely unknown. Second, the most homologous protein to HCAR2 is the same subfamily receptor HCAR3 (GPR109B), exclusively found in humans and higher primates such as chimpanzee.^25^ Notably, HCAR2 shares up to 96% sequence identity with HCAR3, which to some extent increases the difficulty for drug development selectively when targeting the HCAR2 receptor.^26^ A clear example is the niacin and acipimox, which target both HCAR2 and HCAR3, although with a much lower affinity for HCAR3 than for HCAR2.^27^ Last, HCAR2 elicits its physiological responses by coupling primarily to Gi/o proteins to inhibit adenylate cyclase and cyclic AMP signaling. The activation and G protein coupling mechanism underlying HCAR2 are still elusive.

In this study, we employed single-particle cryo-electron microscopy (cryo-EM) to determine the structures of human HCAR2 in complex with heterotrimeric Gi1 protein for the first time: HCAR2 bound to the drugs niacin and acipimox; HCAR2 bound to the highly subtype-specific agonist MK-6892; and HCAR2 in the absence of a ligand (apo) state. Combined with computational modeling, molecular docking, and mutagenesis results, our study provides a structural framework for understanding the ligand recognition and selectivity, receptor activation, and G protein coupling mechanism of HCAR2. More importantly, we believe that these accurate structure templates will accelerate the development of HCAR2-targeting drugs with greater efficacy, higher selectivity, and fewer or no side effects.

## 2 Results

### 2.1 Overall structure of the HCAR2-Gi complex

To investigate the molecular mechanisms of HCAR2 in ligand recognition and signal transduction, we prepared a stable HCAR2-Gi1 complex through co-expression of three subunits of the Gi1 protein and HCAR2 receptor in Sf9 insect cells. Immediately afterward, the HCAR2-Gi1 complex was assembled with scFv16, a Gi-stabilizing antibody, in the absence or presence of an agonist, thus obtaining the cryo-EM density maps of four different complexes with overall resolutions of 3.28 Å (apo), 2.69 Å (niacin), 3.23 Å (acipimox), and 3.25 Å (MK-6892) (Fig. 1a–d). The majority of the side chains of HCAR2 and the Gi1 protein residues were well defined in all obtained complexes, providing accurate models of intermolecular interactions of HCAR2 with the ligand and Gi1 (Supplementary information, Fig. S1–S4). It can clearly be seen that HCAR2 displayed the canonical GPCR topology of a heptahelical transmembrane bundle (7TM), connected by an extracellular N-terminus, three extracellular loops (ECL1–3) and three intracellular loops (ICL1–3). The plotted snake diagram of HCAR2 showed that it also contained an amphipathic helix VIII and a long C-terminus, although their electron densities were not observed in our cryo-EM maps, indicating highly flexible properties of these regions (Supplementary information, Fig. S5).

**Figure 1.**
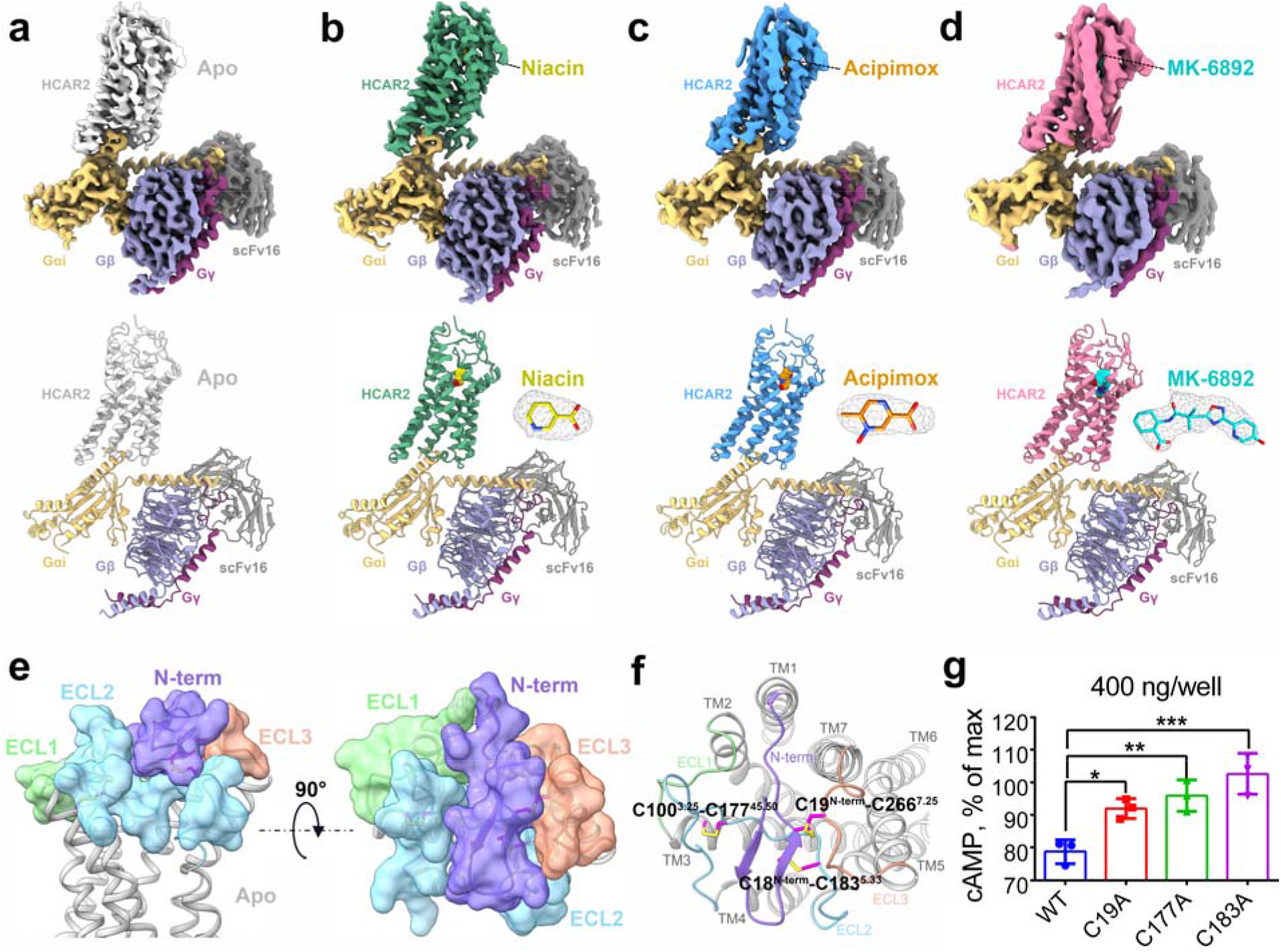
Cryo-EM structures of HCAR2-Gi1 in the apo and niacin-, acipimox-, MK-6892-bound forms. Cryo-EM maps and structural models of HCAR2-Gi1 signaling complex in the absence **(a)** or presence of niacin **(b)**, acipimox **(c)**, and MK-6892 **(d)**. The densities of the agonists (shown as sticks) are depicted as gray meshes. The maps and structural models are colored by subunits. Light gray, apo-HCAR2; forest green, niacin-HCAR2; deep sky blue, acipimox-HCAR2; hot pink, MK-6892-HCAR2; light yellow, Gαi; slate blue, Gβ; dark magenta, Gγ; dark gray, scFv16; yellow, niacin; dark orange, acipimox; cyan, MK-6892. **e** Extracellular architecture of apo-HCAR2 from side and top views. The N-terminal loop (blue purple), ECL1 (light green), ECL2 (sky blue), and ECL3 (coral) are shown as transparent surface presentations and overlaid on the cartoon model of HCAR2 (light gray). **f** Three disulfide bonds (magenta sticks) are formed in the extracellular region of HCAR2. The N-terminal loop (blue purple), ECL1 (light green), ECL2 (sky blue), and ECL3 (coral) are shown as cartoon model. **g** Effect on Gi-mediated cAMP by single-point mutations of C19^N-term^, C177^45.50^, and C183^5.33^ that disrupt the disulfide bonds. The data are presented as mean ± SEM, one-way analysis of variance (ANOVA), \**p* < 0.05, \*\**p* < 0.01, \*\*\**p* < 0.001. The experiments were performed in triplicates.

Despite binding to different ligands, the overall structures of the four active state HCAR2-Gi1 complexes resembled each other, with root-mean-square deviation (RMSD) values of 0.4–1.0□Å for the Cα atoms. A striking feature of HCAR2 was that the ECL2 region (K164–Q187) formed a lid that almost completely capped the extracellular vestibule (Fig. 1e). In particular, a hydrophobic residue F180^ECL2^ in ECL2 was observed to deeply insert into the orthosteric pocket and pack tightly into a local aromatic environment formed by the residues F276^7.35^, F277^7.36^, and F193^5.43^ (Supplementary information, Fig. S6a). This appears to explain the reason for the presence of the apo state of the HCAR2-Gi1 complex. This indicates that HCAR2 does not necessarily require ligand binding to activate the downstream signaling transducers, because its ECL2 may act as a built-in “agonist”. Mutagenesis and cellular functional assays showed that replacing the ECL2 region with a six-residue linker (GGSGGS) or mutating the key residue F180^ECL2^ dramatically reduced the signaling activity of HCAR2 (Supplementary information, Fig. S6b, c). To better understand the structural features of ECL2, sequence and structure alignment of HCAR2 with other reported GPCRs with self-activation, including GPR52, GPR17, and BILF1, were performed.^28–30^ As shown in Supplementary information, Fig. S7a–d, although the overall sequence homology was quite low, all the ECL2 regions occupied the orthosteric binding site to different degrees. The only conserved residue in the ECL2 regions was Cys^45.50^, which formed a disulfide bond with Cys^3.25^ of TM3 in all four GPCRs (Fig. 1f; Supplementary information, Fig. S7e–g). Typically, this disulfide bond (Cys^45.50^-Cys^3.25^) between ECL2 and TM3 presents in most class A GPCRs and plays significant roles in maintaining the architecture of the ligand binding pocket as well as contributing to ligand recognition.^31^ HCAR2 exhibited several distinct features compared with these three GPCRs. The N-terminus C18^N-term^ and C19^N-term^ of HCAR2 formed two extra disulfide bonds with C183^5.33^ of ECL2 and C266^7.25^ of ECL3, respectively (Fig. 1f). Ultimately, under the interactions of a total of three disulfide bonds (C100^3.25^-C177^45.50^, C18^N-term^-C183^5.33^, C19^N-term^-C266^7.25^), HCAR2 displayed a unique extracellular architecture: the ECL2 was closely clamped by ECL1 and ECL3, as well as compressed by the N-terminus from the top (Fig. 1e). Unlike HCAR2, the N-terminus of GPR52 and BILF1 did not form a disulfide bond with ECL2, and only formed one with TM1 and ECL3, respectively (Supplementary information, Fig. S7e, g). As a result, a total of two disulfide bonds were observed in the extracellular structures of GPR52 (C114^3.25^-C193^45.50^, C27^N-term^-C40^1.32^) and BILF1 (C97^3.25^-C174^45.50^, C28^N-term^-C258^ECL3^). In summary, we consider that the multiple disulfide bonds formed further improved the structural stability of ECL2, which played an important role in the self-activation of HCAR2. A single-mutant study of C19^N-term^, C177^45.50^, and C183^5.33^ demonstrated that the disruption of disulfide bonds did not significantly affect HCAR2 expression, but profoundly reduced the constitutive Gi-mediated cAMP signaling (Fig. 1g, Supplementary information, Fig. S8, S9).

### 2.2 Ligand recognition of the HCAR2 receptor

Both niacin (EC_50_ = 0.06–0.25 μM) and acipimox (EC_50_ = 2.6–6 μM) are representative drugs targeting HCAR2, while MK-6892 (EC_50_ = 0.016 μM) is a highly subtype-specific agonist of HCAR2 with a higher affinity (Fig. 2a).^3, 22, 32^ The GTP turnover assay further confirmed Gi1 activation by niacin, acipimox, and MK-6892 in vitro, and their corresponding binding strengths were consistent with their EC_50_ values (Supplementary information, Fig. S10). Compared with the architecture of the apo form, the overall extracellular conformations of the niacin-, acipimox-, and MK-6892-bound HCAR2 complexes were also stabilized by three disulfide bonds, and the orthosteric pocket was capped by ECL2 (Supplementary information, Fig. S11). This seems to be a common feature in the active state of the HCAR2 complex. To determine the differences in ligand recognition of HCAR2, a detailed structure-activity relationship analysis was performed between these three agonist-bound HCAR2 complexes.

**Figure 2.**
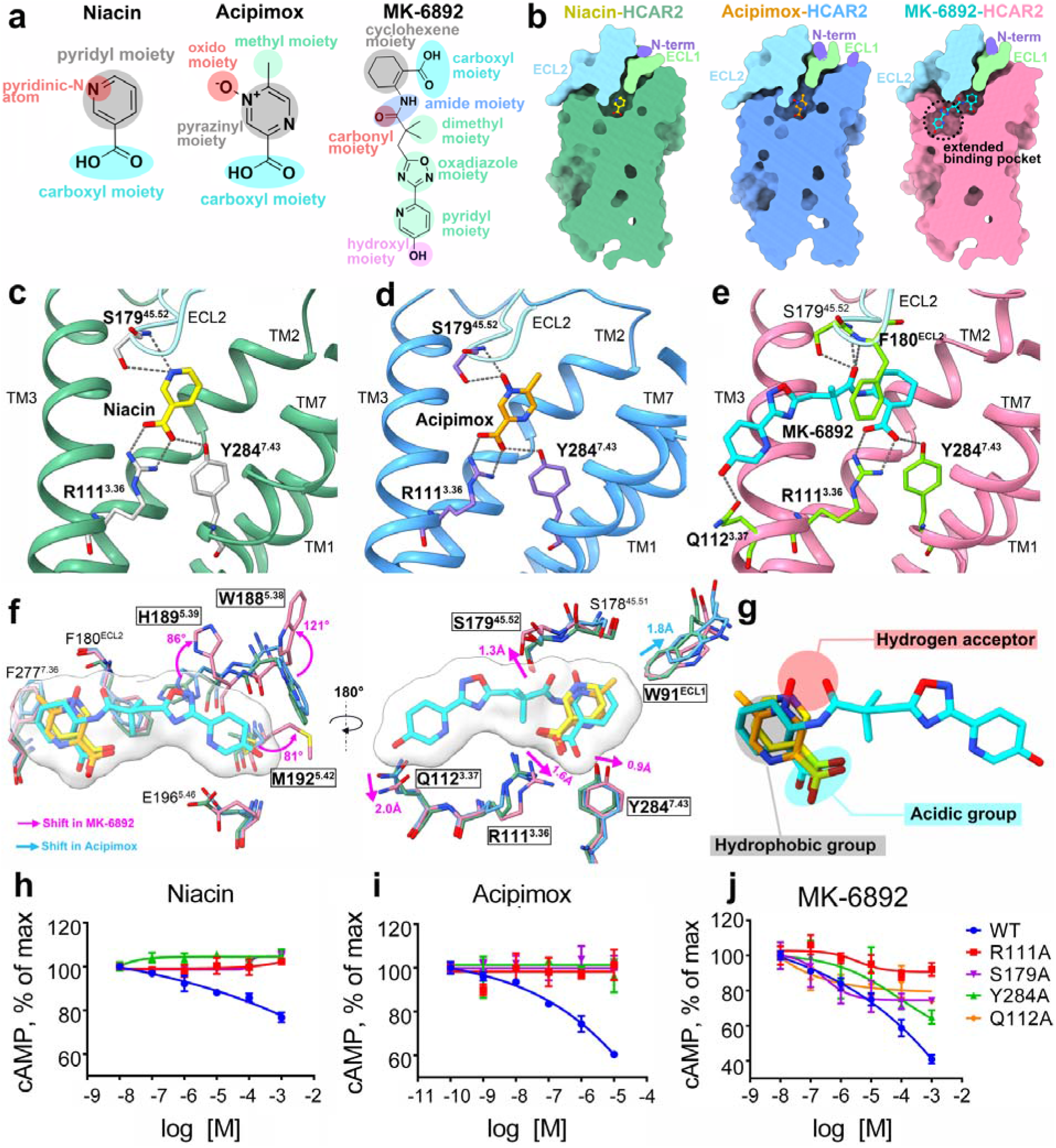
Ligand-binding pocket of active HCAR2 bound to different agonists. **a** Chemical structures of niacin, acipimox, and MK-6892. **b** Vertical cross-sections of the binding pockets of niacin, acipimox, and MK-6892 in HCAR2. **c-e** Detailed interactions of niacin, acipimox and MK-6892 with HCAR2. The polar interactions are indicated by dark gray dashed lines. **f** Superposition of the niacin, acipimox and MK-6892 binding poses, as well as surrounding key residues. **g** General pharmacophore features common to most of the agonists recognized by HCAR2. The structures of HCAR2 and agonists are colored differently. Forest green, niacin-HCAR2; deep sky blue, acipimox-HCAR2; hot pink, MK-6892-HCAR2; blue purple, N-terminal loop; light green, ECL1; sky blue, ECL2; coral, ECL3; yellow, niacin; dark orange, acipimox; cyan, MK-6892; light gray sticks, residues interacting with niacin; medium purple sticks, residues interacting with acipimox; green yellow sticks, residues interacting with MK-6892. **h-j** Effect on Gi-mediated cAMP by single-point mutations of R111^3.36^, Y284^7.43^, S179^45.52^, and Q112^3.37^ that interact with niacin, acipimox, and MK-6892. The data are presented as mean ± SEM. The experiments were performed in triplicates.

When focusing on the structural details of niacin and acipimox complexes, we found that these two ligands adopted a very similar binding pose and were deeply embedded in the orthosteric pocket constituted by TM2, TM3, TM5, TM6, TM7, ECL1, and ECL2 (Fig. 2b, Supplementary information, Fig. S11). In particular, ECL1 and ECL2 were located on the top of the pocket and almost completely isolated the ligands from the extracellular milieu. Further analysis revealed that niacin and acipimox bound to HCAR2 primarily through polar and hydrophobic interactions (Fig. 2c, d, Supplementary information, Fig. S12g, h). Specifically, the positively charged residue R111^3.36^ of HCAR2 was thought to be the most critical residue for binding niacin and acipimox by forming a strong salt bridge with a negatively charged carboxyl group of ligands. The cAMP accumulation assay suggested that the mutation of R111^3.36^ to alanine led to a significant loss of agonistic activity for niacin and acipimox (Fig. 2h, i). Furthermore, the interactions were also stabilized by three pairs of hydrogen bonds: the carboxyl moiety of niacin (or acipimox) with the side chain of Y284^7.43^; and the pyridinic-N atom of niacin (or the oxido moiety of acipimox) with the side chain and backbone of S179^45.52^ (Fig. 2c, d). Consistently, mutation in either Y284^7.43^ or S179^45.52^ markedly reduced ligand activity (Fig. 2h, i). In addition, a hydrophobic environment in the orthosteric pocket was necessary for HCAR2 activation. This was evident from the featured aromatic ring of niacin and acipimox that was surrounded by a series of hydrophobic residues (L83^2.60^, W91^ECL1^, M103^3.28^, L104^3.29^, L107^3.32^, F180^ECL2^, F276^7.35^, F277^7.36^, and L280^7.39^), thus establishing the hydrophobic interactions (Supplementary information, Fig. S12a, b). Overall, the hydrophilic, hydrophobic, and charged properties of niacin and acipimox matched quite well with those of the allosteric pocket: the upper portion of the binding site (toward the extracellular side) was largely hydrophobic, while the bottom portion (intracellular side) was hydrophilic and charged (Supplementary information, Fig. S12d, e). It was still of note that a subtle conformational difference existed between the allosteric pockets of niacin and acipimox complexes. Since acipimox has an additional methyl moiety at the 5-position of the pyrazine ring, W91^ECL1^ was observed to move 1.8□Å toward the top of the pocket so as to accommodate this group (Fig. 2f).

Unlike niacin and acipimox, the ligand-binding pocket formed by bulky MK-6892 was comprised of two subpockets: a canonical orthosteric binding pocket and an extended binding pocket (Fig. 2b). Similar to niacin and acipimox, the carboxyl, cyclohexene, and amide moieties of MK-6892 primarily occupied the orthosteric pocket and formed similar interactions with R111^3.36^, Y284^7.43^, S179^45.52^, and the surrounding hydrophobic residues (Fig. 2e). The only difference was that the carbonyl oxygen of MK-6892 introduced an additional hydrogen bond to the backbone nitrogen of F180^ECL2^, which was absent in the niacin-, and acipimox-bound HCAR2 complexes. This was attributed to the greater interatomic distance of the oxygen and nitrogen in the niacin (3.9□Å) and acipimox (3.7□Å) complexes, while it was only 2.9□Å in the MK-6892 complex (Supplementary information, Fig. S12g–i). Furthermore, the dimethyl, oxadiazole, and pyridyl moieties of MK-6892 were mainly positioned on the extended binding pocket formed by TM3, TM4, and TM5, and made hydrophobic interactions with several residues around them, including A108^3.33^, L158^4.56^, L162^4.60^, A191^5.41^, M192^5.42^, and F193^5.43^ (Supplementary information, Fig. S12c, f). Meanwhile, the top hydroxyl group of MK-6892 formed a hydrogen bond with Q112^3.37^. The mutations of R111^3.36^, Y284^7.43^, S179^45.52^, and Q112^3.37^ to alanine significantly impaired the MK-6892 activity, further confirming their important roles in MK-6892 binding (Fig. 2j). Following the above analysis, we speculated that these extra interactions, particularly those established in the extended binding pocket, contributed to the high binding affinity of MK-6892. Of course, to accommodate such interaction patterns, several minor changes in chemical shifts were observed in the key residues of the MK-6892 complex. For example, compared with the niacin and acipimox complexes, the side chains of R111^3.36^, Q112^3.37^, S179^45.52^, and Y284^7.43^ were found to move 1.6□Å, 2.0□Å, 1.3Å, and 0.9□Å, respectively, to better make polar interactions with the bulky MK-6892 (Fig. 2f). But more importantly, how did the extended binding pocket form in the MK-6892-HCAR2 complex? We noted that the oxadiazole and pyridyl groups of MK-6892 forced the side chains of H189^5.39^ and M192^5.42^ to rotate about 86° and 81°, respectively, thereby avoiding to clash with the ligand (Fig. 2f). Immediately afterward, the rotation of M192^5.42^ occupied the position initially occupied by W188^5.38^ and caused it to rotate upward about 121°. Eventually, these large conformational changes of W188^5.38^, H189^5.39^, and M192^5.42^ in the side chain orientations together led to the extended binding pocket formation.

### 2.3 Pharmacophore features of the HCAR2 agonist

In terms of chemical structures, niacin and acipimox share similar features, including a carboxyl moiety, an aromatic ring (pyridyl moiety of niacin and pyrazinyl moiety of acipimox), and an electron-rich moiety (pyridinic-N atom of niacin and oxido moiety of acipimox) (Fig. 2a). In comparison, MK-6892 is much more complicated. Structural analysis suggested that MK-6892 not only exhibited the shared three structural features of niacin and acipimox, but also had several unique groups, including dimethyl, oxadiazole, pyridyl, and hydroxy moieties (Fig. 2a). The previous section had provided important insights into the key roles of different moieties in niacin, acipimox, and MK-6892 for HCAR2 activation. Therefore, what are the structural features of ligand recognition of HCAR2?

On the basis of the acipimox-bound HCAR2 structure, we investigated the interactions of HCAR2 with more HCAR2 agonists, including endogenous ligands (butyrate and β-OHB), important drugs (MMF and acifran), and 3-pyridineacetic acid, through molecular docking (Supplementary information, Fig. S13a).^33^ The docking pose of acipimox reproduced the crystallographic pose well, increasing the accuracy and reliability of the docking results (Supplementary information, Fig. S13b). The predicted binding modes suggested that all agonists were bound nicely in the orthosteric pocket. Except for butyrate, all other agonists mainly formed polar interactions with R111^3.36^, Y284^7.43^, and S179^45.52^, which were consistent with those of niacin and acipimox (Supplementary information, Fig. S13c–g). It is well known that both butyrate and β-OHB are endogenous ligands of HCAR2, but butyrate lacks one hydroxyl group in its structure, making it unable to form a hydrogen bond with S179^45.52^. This result provided a good explanation for why β-OHB has a higher potency for HCAR2 than butyrate.^2, 6^ Additionally, there were still hydrophobic interactions between these agonists and surrounding hydrophobic residues. The difference was that acifran and 3-pyridineacetic acid interacted mainly through a rigid aromatic ring, like that of niacin, acipimox, and MK-6892, whereas butyrate, β-OHB, and MMF interacted primarily via aliphatic chains. Combining the cryo-EM structures and docking results, we summarized the general pharmacophore features that may be common to most of the agonists recognized by HCAR2: an acidic group (contributes to the salt bridge and hydrogen bond with basic R111^3^.^36^ and Y284^7.43^), a hydrogen acceptor (contributes to the hydrogen bond with S179^45.52^), and a hydrophobic aliphatic or aromatic group (contributes to the hydrophobic interactions) (Fig. 2g). Among them, the negatively charged acidic group was considered to be the most important and essential factor for HCAR2 activation. This is because previous studies suggested that if the carboxyl group of niacin was replaced with an amide group, the produced nicotinamide was no longer active toward HCAR2.^34^

### 2.4 Ligand selectivity between HCAR2 and HCAR3 receptors

Of the entire hydroxycarboxylic acid receptor family, both HCAR2 and HCAR3 share relatively low homology with HCAR1, because their amino acid sequence identities are only 48.9% and 47.0%, respectively. By comparison, there is up to 96% sequence identity between HCAR2 and HCAR3, which differ by only 15 amino acid residues in most of their domains, including the N-terminus, TM1–7, ECL1–3, and ICL1–3 (Supplementary information, Fig. S14). Specifically, eight amino acids cluster in the TM1–7; one amino acid clusters in the ECL1; and six amino acids cluster in the ECL2 (Fig. 3a). Moreover, HCAR3 has an extended C-terminus containing 24 additional amino acids. Given that the two receptors are highly homologous, this allowed us to model the HCAR3 structure relatively accurately based on our resolved HCAR2 structures. Through analysis of conformational differences, we decided to explore the possible reasons why some agonists, such as niacin and acipimox, can bind to both HCAR2 and HCAR3, but display higher selectivity for HCAR2.^13^ These findings are especially important for the development of more selective HCAR2 - targeting drugs.

**Figure 3.**
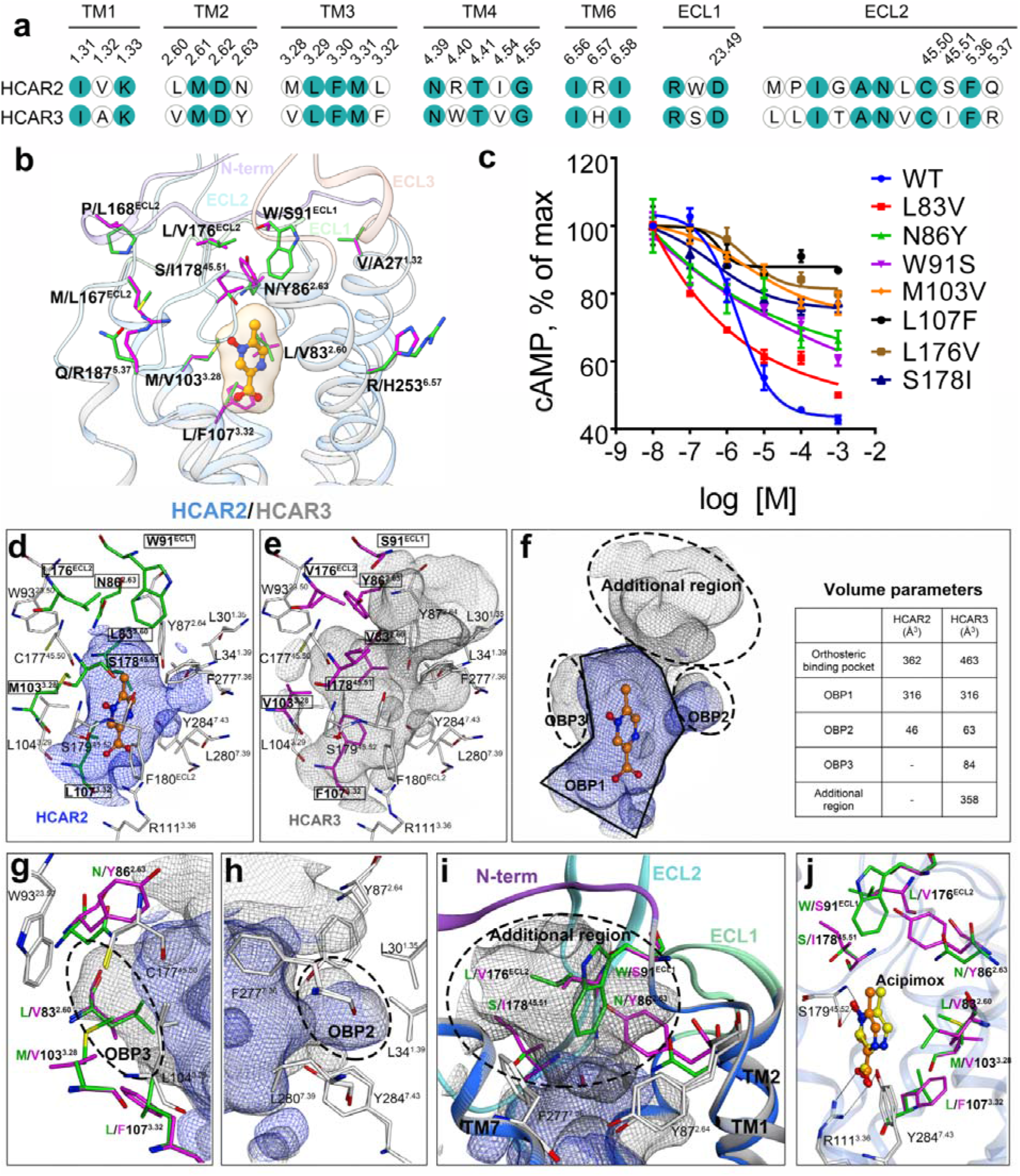
Structural basis for ligand selectivity between HCAR2 and HCAR3. **a** Sequence alignment of residues in HCAR2 and HCAR3. The conserved residues are highlighted in solid dark green circles. **b** Superposition of the 15 different residues. The N-terminal loop (blue purple), ECL1 (light green), ECL2 (sky blue), and ECL3 (coral) in HCAR2 (deep sky blue) and HCAR3 (gray) are overlaid in cartoon representation. The 15 different residues in the HCAR2 (green), HCAR3 (magenta), and acipimox (dark orange) are shown in stick representation. **c** Effect on Gi-mediated cAMP by single-point mutations of L83^2.60^, N86^2.63^, M103^3.28^, L107^3.32^, W91^ECL1^, L176^ECL2^, and S178^45.51^ related to the formation of the orthosteric binding pocket. The data are presented as mean ± SEM. The experiments were performed in triplicates. **d-i** The orthosteric binding pocket of HCAR2 (blue) and HCAR3 (dark gray) are overlaid. The different residues in HCAR2 (green) and HCAR3 (magenta), as well as the surrounding conserved residues (white) are shown in stick representation. The N-terminal loop (blue purple), ECL1 (light green), ECL2 (sky blue), and ECL3 (coral) in HCAR2 (deep sky blue) and HCAR3 (gray) are shown in cartoon representation. **j** Superposition of acipimox between the crystal structure (dark orange) and docking conformation (yellow). The different residues in HCAR2 (green) and HCAR3 (magenta) are shown in stick representation.

The homology model for HCAR3 was constructed using the acipimox-bound HCAR2 complex as a template with the SWISS-MODEL online server. Then, we performed fine mapping of the 15 residues differing in the HCAR2 and HCAR3 structures (Fig. 3b, Supplementary information, Fig. S15). With the exception of three residues located at positions 142, 156, and 173, the remaining 12 residues were found around the coordinates of the bound ligand. In particular, 7 of 12 residues, located at positions 83, 86, 91, 103, 107, 176, and 178, were directly related to the formation of the orthosteric binding pocket (Fig. 3d, e). The differences in these seven residues not only rendered the pocket volume ~100 Å^3^ larger in HCAR3 than in HCAR2, but also generated an additional region (~358 Å^3^) at the top of the pocket. For better comparison, we overlaid their orthosteric binding pockets and subdivided them into three parts, which were defined as OBP1, OBP2, and OBP3 (Fig. 3f). As the main cavity that accommodated the ligand, the OBP1 regions of HCAR2 and HCAR3 were well overlapped, and their volumes were ~316 Å^3^. The OBP2 regions did not differ significantly, and had volumes of ~46 Å^3^ (HCAR2) and ~63 Å^3^ (HCAR3) (Fig. 3h). This is primarily because the residues comprising OBP2, including L30^1.35^, L34^1.39^, Y87^2.64^, F277^7.36^, L280^7.39^, and Y284^7.43^, were all highly conserved. In contrast, the most remarkable difference was the extra cavity (OBP3 region) with ~84 Å^3^ in HCAR3, which was absent in HCAR2 (Fig. 3g). There were seven residues associated with the OBP3 formation, but four differed among them: L83^2.60^, N86^2.63^, M103^3.28^, and L107^3.32^ were in HCAR2, while V83^2.60^, Y86^2.63^, V103^3.28^, and F107^3.32^ were in HCAR3. Mutagenesis study revealed that the substitution of the four residues in HCAR2 with the corresponding residues in HCAR3 led to significant mitigation of downstream Gαi stimulation by acipimox in HCAR2 (Fig. 3c). Another difference of note was the additional region of HCAR3, the formation of which was related to the replacement of the residues at positions 86, 91, 176, and 178 (Fig. 3i). Particularly, the bulky residue W91^ECL1^ in the HCAR2 was positioned on the top of the orthosteric pocket like a lid, whereas it was substituted with a smaller residue, S91^ECL1^, in HCAR3, thus producing a large additional region composed of the N-terminus, ECL1, ECL2, TM1, TM2, and TM7. Consistently, single mutation of W91^ECL1^, L176^ECL2^, and S178^45.51^ in HCAR2 to S91^ECL1^, V176^ECL2^, and I178^45.51^ in HCAR3 also markedly reduced the acipimox activity (Fig. 3c). Subsequently, we docked acipimox into the orthosteric pocket of HCAR3 and selected the highest ranked pose as the probable binding conformation (Fig. 3j). The key amino acid residues R111^3.36^, Y284^7.43^, and S179^45.52^ were conserved in the HCAR3 sequence, and thus the predicted binding mode of acipimox in HCAR3 resembled that in the HCAR2 crystal structure overall. However, the conformation of acipimox in HCAR3 deflected with an angle (~23°) and the corresponding binding free energy (−31.82 kcal/mol) was higher than that of acipimox with HCAR2 (−39.39 kcal/mol), indicating a relatively weak affinity toward HCAR3. In view of these results, we speculated that the differences in the pocket volume and shape of the two receptors, chiefly contributed by residues at positions 83, 86, 91, 103, 107, 176, and 178, had a significant influence on the agonist selectivity between HCAR2 and HCAR3.

### 2.5 Activation of the HCAR2 receptor

The four active state HCAR2-Gi1 complexes enabled us to explore the activation mechanism of HCAR2. As can be clearly seen from Fig. 4a, the overall structure of the TM1–7 regions and several canonical activation-related motifs, such as P^5.50^-I^3.40^-F^6.44^, N/D^7.49^P^7.50^xxY^7.53^, and E/D^3.49^R^3.50^Y^3.51^ (where x is any residue), are highly superimposed in the apo and agonist-bound HCAR2 structures, suggesting a shared activation mechanism (Supplementary information, Fig. S16a-c). Typically, the activation motion of most class A GPCRs is triggered by a conserved residue W^6.48^, which serves as the “toggle switch” in TM6.^35^ Afterward, the movement of W^6.48^ gives rise to rearrangement of the conserved P^5.50^-I^3.40^-F^6.44^ motif and leads to outward movement of TM6.^36^ When the activation signal is propagated through the conserved N/D^7.49^P^7.50^xxY^7.53^ motif to the bottom E/D^3.49^R^3.50^Y^3.51^ motif, TM6 moves further outward to accommodate the binding of G proteins.^37^ Due to the lack of an inactive structure, the detailed conformational changes that HCAR2 experienced are still unknown. But like most class A GPCRs, these important motifs mentioned above were completely conserved in HCAR2, as the corresponding positions were P200^5.50^-I115^3.40^-F240^6.44^, D290^7.49^P291^7.50^xxY294^7.53^, and D124^3.49^R125^3.50^Y126^3.51^ (Supplementary information, Fig. S16a-c). Notably, in HCAR2, the conserved W^6.48^ was replaced by F244^6.48^, which established extensive aromatic and hydrophobic interactions with surrounding residues, including the P200^5.50^-I115^3.40^-F240^6.44^ motif and F197^5.47^ (Supplementary information, Fig. S16a). To our knowledge, this change is common and present in many δ-branch class A GPCRs (e.g., protease-activated receptors (PARs) and cysteinyl leukotriene receptors (CysLTRs)), because many of them have F^6.48^ instead of W^6.48^ at this position.^38, 39^ Through the above analysis, we considered that HCAR2 also followed the same general activation processes seen in other class A GPCRs.

**Figure 4.**
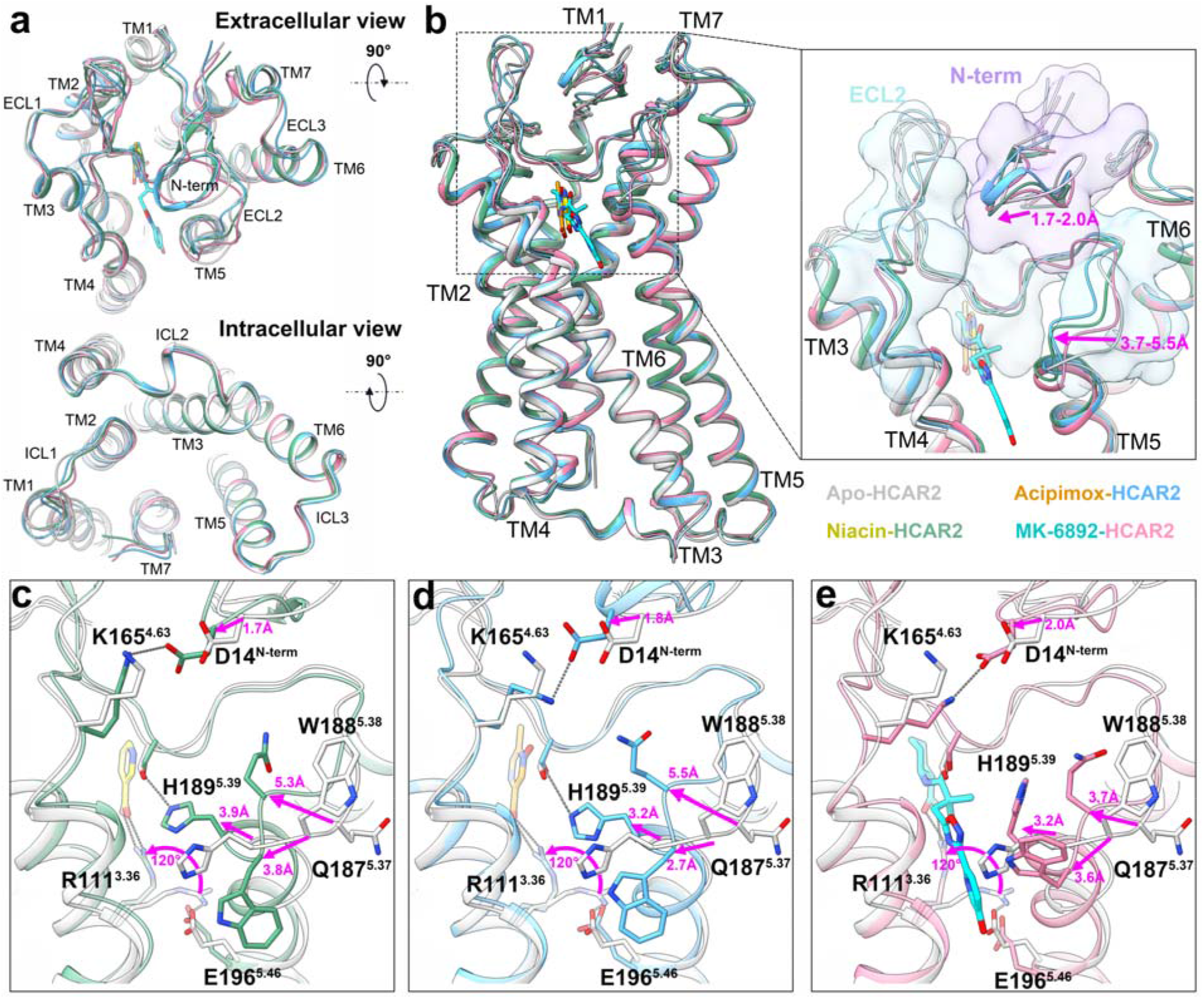
Structural comparison of HCAR2 in four active states. **a** Extracellular and intracellular views of the comparison of the apo and agonist-bound HCAR2 structures. **b** Structural differences of the N-terminus and ECL2 in the four active state HCAR2 receptors. Pairwise comparisons of the apo state versus niacin-**(c)**, acipimox-**(d)**, and MK-6892-bound **(e)** forms. The structures of HCAR2 and agonists are colored differently. Light gray, apo-HCAR2; forest green, niacin-HCAR2; deep sky blue, acipimox-HCAR2; hot pink, MK-6892-HCAR2; blue purple, N-terminal loop; sky blue, ECL2; yellow, niacin; dark orange, acipimox; cyan, MK-6892; magenta arrow, shift with respect to the apo state; dark gray dashed lines, polar interactions.

Interestingly, despite that the four active state complexes had similar activation patterns, the ligand binding seemed to make the HCAR2 structure tighter and more stable, which was predominately reflected in the N-terminus and ECL2. We performed pairwise comparisons for the apo state versus niacin-, acipimox-, and MK-6892-bound forms. Compared with the apo state, the N-terminus and ECL2 in the agonist-bound HCAR2 structures were observed to move 1.7–2.0 Å and 3.7-5.5 Å, respectively (Fig. 4b). Specifically, the movement of the N-terminus was mainly mediated by D14^N-term^, which closed the distance between the N-terminus and ECL2 by forming a salt bridge with K165^4.63^ (Fig. 4c-e). Additionally, in the apo state, the residue R111^3.36^ formed an “ionic lock” with E196^5.46^. However, the addition of agonist disrupted this interaction and induced R111^3.36^ to rotate upward ~120° to establish a salt bridge with an acidic group of the agonist. Meanwhile, a large structural rearrangement occurred in Q187^5.37^, W188^5.38^, and H189^5.39^ of ECL2, which moved 3.7–5.5 Å, 2.7–3.8 Å, and 3.2–3.9 Å, respectively, and rotated at different degrees. Together these changes eventually led to the formation of multiple polar interactions between the agonist and R111^3.36^, Y284^7.43^, and S179^45.52^, as we described previously in niacin-, acipimox-, and MK-6892-bound complexes (Fig. 2c–e). Thus, the ligand binding, to some extent, increased the structural stability of HCAR2 by tethering TM3, TM7, and ECL2 together. It was worth noting that the conformational change of residue H189^5.39^ had been regarded as important for the formation of the orthosteric binding pocket in the MK-6892-bound form (Fig. 2f). Here, we further observed that in the niacin- and acipimox-bound forms, H189^5.39^ extended to the ligand binding orientation and formed a hydrogen bond with S179^45.52^, making the ECL2 conformation more stable (Fig. 4c, d).

### 2.6 Interfaces between the HCAR2 receptor and Gi1

The complex structures of HCAR2-Gi1 with or without agonists showed almost the same G protein coupling interface. As depicted in Fig. 5a, the interactions between HCAR2 and Gi1 were mainly mediated by the α5 helix of the Gαi subunit and the receptor cores comprised by TM1–3, TM5, TM6, ICL2, and ICL3. The αN helix of the Gαi subunit here had almost no direct interaction with the receptor. Similar to many other Gi-bound class A GPCRs, the C-terminus of the α5 helix was amphipathic and inserted into the cytoplasmic cavity of HCAR2 by forming extensive hydrophobic and electrostatic interactions, as well as hydrogen bonds (Fig. 5c-f). Three large hydrophobic side chains L353, L348, and I344 of the α5 helix embedded in a hydrophobic groove of HCAR2 formed by TM3, TM5/6, and ICL3 residues (TM3: V129^3.54^; TM5: I211^5.61^ and L215^5.65^; TM6: I226^6.30^, A229^6.33^, F232^6.36^, and I233^6.37^; and ICL3: M220^ICL3^), forming extensive hydrophobic interactions. Electrostatic interactions were also crucial for the stabilization of the binding mode between HCAR2 and Gi1. HCAR2 displayed highly positive charges at the cytoplasmic ends of TM2 and ICL3, contributed by K60^2.37^ and R218^ICL3^. Correspondingly, the α5 helix C-terminus was highly negatively charged, contributed by D350 and D341, and ultimately established salt bridges between α5 and HCAR2. In addition, the residues R128^3.53^, H133^34.51^, and S298^8.47^ of HCAR2 formed well-defined hydrogen bonds with N347, T340, and G352 of the α5 helix, respectively. We then aligned the HCAR2-Gi1 complex with several canonical class A GPCR complexes, such as rhodopsin, μ-opioid receptor (μOR), and cannabinoid receptor 1 (CB1), as well as self-activated GPR17, and observed that the major differences occurred in the relative positions and orientations of the α5 and αN helices, as well as the shift of TM6 (Fig. 5b).^30, 40–42^ For example, HCAR2 had a much less pronounced outward movement at the cytoplasmic end of TM6 compared to the representative GPCRs. Relative to the HCAR2-Gi1 complex, the α5 and αN helices rotated ~3.5 Å and ~7.6 Å in rhodopsin-Gi1; ~4.9 Å and ~8.1 Å in μOR-Gi1; ~1.4 Å and ~3.3 Å in CB1-Gi1; and ~2.7 Å and ~13.1 Å in GPR17-Gi1, respectively (Supplementary information, Fig. S17a-d). The shifts of Gi1 coupling were most likely attributed to the differences in receptor structures. Together, all these findings clarified the Gi1 coupling features of HCAR2 and provided a greater understanding of the G protein coupling mechanism.

**Figure 5.**
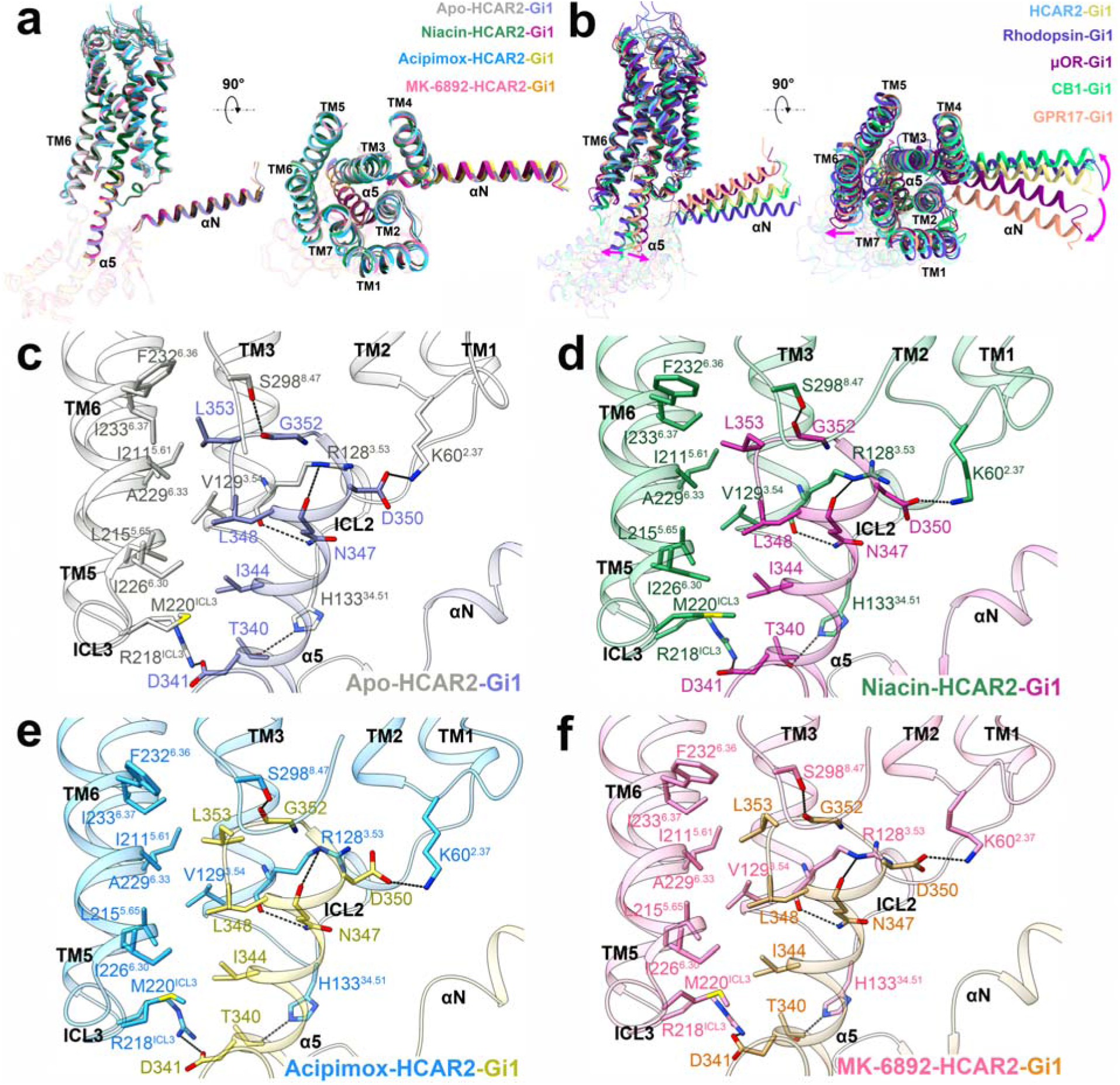
Analysis of the HCAR2-Gi1 interface and comparison with class A GPCRs. **a** Comparison of the HCAR2-Gi1 complex with or without an agonist. **b** Superimposition of the receptor G protein coupling interfaces for HCAR2-Gi1, rhodopsin-Gi1 (PDB: 6CMO, slate blue), μOR-Gi1 (PDB: 6DDE, dark magenta), CB1-Gi1 (PDB: 6N4B, turquoise), and GPR17-Gi1 (PDB: 7Y89, dark salmon). **c-f** Interactions of HCAR2 with the α5 helix of Gαi. The structures of HCAR2 and Gi1 are colored differently. Light gray and purple, apo-HCAR2-Gi1; forest green and plum, niacin-HCAR2-Gi1; deep sky blue and light yellow, acipimox-HCAR2-Gi1; hot pink and orange, MK-6892-HCAR2-Gi1; dark gray dashed lines, polar interactions; magenta arrow, shift with respect to HCAR2-Gi1.

## 3 Discussion

In recent decades, a series of HCAR2 agonists have been successfully discovered and four of them, including niacin, acipimox, acifran, and MMF, have been approved for clinical treatment of cardiovascular and neurological disorders, such as dyslipidemia, atherosclerosis, and relapsing multiple sclerosis.^13^ Despite the favorable clinical efficacy, all four drugs can cause the unwanted side effect of cutaneous flushing.^43, 44^ It is known that flushing is a cutaneous vasodilation accompanied by a burning sensation mainly affecting the upper body and face.^4^ There is good evidence that cutaneous flushing is associated with the activation of HCAR2 at Langerhans cells and keratinocytes, as well as the subsequent release of vasodilatory prostaglandins.^43^ In light of this, some highly subtype-specific HCAR2 agonists with fewer side effects have been developed, especially MK-6892. In vivo rat and dog experiments demonstrated that MK-6892 displayed excellent therapeutic index compared to niacin in free fatty acid reduction, but significantly relieved the flushing effect.^22^ Unfortunately, however, all the structures of hydroxycarboxylic acid receptors, HCAR1–3, have not been resolved, and it is still unknown what the similarities and differences are of the recognition mechanisms and binding modes between the four approved drugs and MK-6892.

In the present study, we first reported four cryo-EM structures of HCAR2-Gi1 complexes in the apo and niacin-, acipimox-, MK-6892-bound forms. For the apo-HCAR2 structure, we showed that ECL2 operated as a built-in “agonist” to activate HCAR2 without requiring ligand binding. Combining the niacin- and acipimox-bound HCAR2 structures, as well as the molecular docking results, we found that many agonists, including butyrate, β-OHB, niacin, acipimox, acifran, and MMF, all adopted similar binding poses and bound to HCAR2 by directly interacting with three key residues, R111^3.36^, S179^45.52^, and Y284^7.43^, in the orthosteric binding pocket. Among them, the salt bridge between an acidic group and basic R111^3.36^ was considered as the most important interaction, because replacement of the carboxyl group with an amide group could not activate HCAR2.^34^ Based on this, the detailed recognition mechanisms of HCAR2 for endogenous ligands and four important drugs were revealed, which are critical for understanding how these agonists exert their anti-lipolytic and anti-inflammatory functions. More importantly, the general pharmacophore features that may fit most of the agonists recognized by HCAR2 were summarized.

Compared with niacin and acipimox, highly subtype-specific MK-6892 had several unique groups, resulting in a larger and more complex chemical structure. To accommodate the bulky MK-6892, substantial conformation changes of W188^5.38^, H189^5.39^, and M192^5.42^ led to the formation of an extended binding pocket in the MK-6892-HCAR2 complex. As a result, MK-6892 not only formed similar interactions with R111^3.36^, Y284^7.43^, and S179^45.52^ as niacin and acipimox in the orthosteric pocket, but also introduced additional polar and hydrophobic interactions in the extended binding pocket. On one hand, this well explained the reason for the higher affinity of HCAR2 for MK-6892 than for niacin and acipimox.^22^ On the other hand, we speculated that the differences in the binding modes of MK-6892 and of niacin and acipimox might differentially activate downstream signaling pathways, thus selectively eliciting the therapeutic, anti-lipolytic pathway, while avoiding the activation of the flush-inducing pathway. In fact, the concept that some agonists may selectively activate only a subset of signal effectors via a common receptor has been confirmed in a number of GPCRs.^44, 45^

As a close relative of HCAR2, HCAR3 is considered to be the result of a recent gene duplication present in humans and higher primates, such as chimpanzee.^1, 46^ Unlike HCAR2, the natural ligand of HCAR3 is 3-hydroxyoctanoic acid, which exerts anti-lipolytic activity in the human body.^46^ Interestingly, HCAR2 is not activated by 3-hydroxyoctanoic acid, and the HCAR2 natural ligands β-OHB and butyrate have no activity toward HCAR3 as well.^47^ Furthermore, some agonists, such as niacin and acipimox, can bind to both receptors, but display higher selectivity for HCAR2.^13^ To explore the ligand selectivity differences between HCAR2 and HCAR3, we constructed a model of HCAR3 relatively accurately using the acipimox-bound HCAR2 complex as a template. Structural alignment suggested that although only 15 differential residues were observed in the major domains of HCAR2 and HCAR3, seven of them, at positions 83, 86, 91, 103, 107, 176, and 178, significantly affected the volume and shape of the orthosteric binding pockets: (1) the pocket volume of HCAR3 was larger than that of HCAR2; (2) an additional region was generated on the top of the HCAR3 pocket. After docking acipimox into the orthosteric pocket of HCAR3, the overall binding mode of acipimox in HCAR3 was similar to that in HCAR2. This was mainly because the key residues R111^3.36^, Y284^7.43^, and S179^45.52^ were conserved and well overlapped in the two receptors. Nevertheless, a higher binding free energy in HCAR3 indicated that acipimox had a higher affinity for HCAR2 than HCAR3, which meant that the pocket differences indeed affected agonist selectivity. We conjectured that the smaller pocket volume of HCAR2 might be more favorable for precise positioning and binding of acipimox to the surrounding residues, thus forming stable interactions. Our results were also confirmed by the study of Ahmed et al., in which the residues at positions 86, 103, and 107 were considered to be critically involved in forming the selective binding site in HCAR3.^47^ To get more details on the precise interactions between ligands and HCAR3, the cryo-EM structures of agonist-bound HCAR3 are in progress. Overall, our structural analysis provides a deep understanding of the ligand recognition, selectivity, activation, and G protein coupling mechanism of HCAR2, which is important for the design of HCAR2-targeting drugs with greater efficacy, higher selectivity, and fewer or no side effects.

## 4 Materials and Methods

### 4.1 Expression and purification of the HCAR2-Gi1 complex

Wild-type human HCAR2 was cloned into pFastBac vector (Gibco) with N-terminal hemagglutinin (HA) signal sequence, Flag tag, and HRV 3C protease site, as well as a C-terminal His tag. Dominant-negative Gαi1 (DNGαi1) with mutations (G203A and A326S) was constructed in the same manner as HCAR2. The pFastBac Dual vector (Gibco) was used to construct the Gβ_1_γ_2_ expression vector. The co-expression of HCAR2, DNGαi1, and Gβ_1_γ_2_ proteins was achieved using the Bac-to-Bac baculovirus expression system in *Spodoptera frugiperda* Sf9 cells (Invitrogen). Cells were grown in suspension at 27°C to a density of 4□×□10^6^ cells ml^1^ and infected with virus at a ratio of 10:10:1 (HCAR2: DNGαi1: Gβ1γ_2_). After 48 h of infection, the cells were collected by centrifugation and then stored at −80°C until use.

To obtain the HCAR2-Gi1 complex, cell pellets were thawed and suspended in lysis buffer (10 mM HEPES, pH 7.5, 0.5 mM EDTA) and supplemented with 50 μM niacin (MCE HY-B0143), acipimox (MCE HY-B0283), MK-6892 (MCE HY-10680), or without an agonist. Then, the sample was rotated at 4°C for 60 min to induce HCAR2-Gi1 complex formation. A dounce homogenizer was used to homogenize and collect cell membranes in a solubilization buffer [20 mM HEPES (pH 7.5), 100 mM NaCl, 50 μM agonist, 10% glycerol, 1% (w/v) n-Dodecyl-b-D-maltoside (DDM), 0.1% (w/v) cholesteryl hemisuccinate (CHS), 0.2 μg ml^-1^ leupeptin, 100 μg ml^-1^ benzamidine, 10 mM MgCl_2_, 5 mM CaCl_2_, 1 mM MnCl_2_, 100 μU ml^-1^ lambda phosphatase (NEB), and 25 μU ml^-1^ apyrase (NEB)]. After incubating at 4°C for 2 h, the supernatant was collected by centrifugation and then incubated with anti-Flag M1 antibody affinity resin at 4°C for 1 h. The M1 resin was washed with wash buffer [20 mM HEPES (pH 7.5), 100 mM NaCl, 50 μM agonist, 0.1% DDM, 0.01% CHS, and 2 mM CaCl_2_]. The buffer was changed from DDM to lauryl maltose neopentyl glycol (LMNG) using a stepwise process. Afterward, the M1 resin was washed with the LMNG buffer [20 mM HEPES (pH 7.5), 100 mM NaCl, 50 μM agonist, 0.01% (w/v) LMNG, 0.001% CHS, and 2 mM CaCl_2_]. The HCAR2-Gi1 complex was eluted with elution buffer [20 mM HEPES (pH 7.5), 100 mM NaCl, 50 μM agonist, 0.00075% LMNG, 0.00025% (w/v) glycol-diosgenin (GDN), 0.0001% CHS, 5 mM EDTA, and 200 μg ml^-1^ synthesized Flag peptide]. The eluted protein was concentrated and incubated with the antibody fragment scFv16 for 2 h on ice at a molar ratio of 1:1.5.^48^ A Superdex 200 Increase 10/300 column (GE Healthcare) was pre-equilibrated with buffer [20 mM HEPES (pH 7.5), 100 mM NaCl, 0.00075% LMNG, 0.00025% GDN, 0.0001% CHS, and 50 μM agonist] and then used to further purify the complex. The obtained pure HCAR2-Gi1-scFv16 complex was concentrated with an ultrafiltration tube and flash frozen in liquid nitrogen until further use.

### 4.2 GTPase GLO assay

To perform the GTPase-Glo assay, the HCAR2 protein was purified as described above. Then we initiated the GTPase reaction by mixing Gi1 and HCAR2 in 5 μl of reaction buffer [20 mM HEPES (pH 7.5), 100 mM NaCl, 0.02% LMNG, 1 mM MgCl2, 5 μM GTP, 5 μM GDP, with or without 50 μM test agonist] in a 384-well plate. The final concentration of HCAR2 and Gi1 were 4 μM and 0.5 μM, respectively. Gi1 alone was set as a reference in every independent experiment. At room temperature (22–25°C), the GTPase reaction was incubated for 2□h. Then 5 μl of reconstituted 1xGTPase-Glo reagent (Promega) was added, mixed briefly, and incubated with shaking for 30 min to convert the remaining GTP into ATP. Afterward, to convert the ATP into luminescent signals, we added 10 μl of detection reagent (Promega) to the system, which was incubated in the 384-well plate for 5–10 min at room temperature. The Multimode Plate Reader (PerkinElmer EnVision 2105) luminescence counter was used to quantify the luminescence intensity. Data were analyzed using GraphPad Prism 9.0.

### 4.3 Cryo-grid preparation and EM data collection

To prepare the cryo-EM sample, the 100 Holey Carbon film (Au, 300 mesh, N1-C14nAu30-01) was pre-discharged with Tergeo-EM plasma cleaner. Then, 3 μl of the purified HCAR2-Gi1-scFv16 complex was applied to the grid. At 10°C and 100% humidity, the sample was incubated for 3 s and blotted for 2 s using the freezing plunger Vitrobot I (Thermo Fisher Scientific, USA). Grids were quickly frozen in liquid ethane cooled by liquid nitrogen and stored in liquid nitrogen until checked. We used the 300 kV Titan Krios Gi3 microscope (Thermo Fisher Scientific FEI, the Kobillka CryoEM Center of the Chinese University of Hong Kong, Shenzhen) to check the grids and collect the cryo-EM data of the HCAR2-Gi1-scFv16 complex. The Gatan K3 BioQuantum camera at a magnification of 105,000 was used to record movies, and the pixel size was 0.83–0.85 Å. We used the GIF-quantum energy filter (Gatan, USA) to exclude the inelastically scattered electrons. The slit width of the filter was set to 20□eV. The movie stacks were automatically acquired with the defocus range from −1.1 to −2.0 μm. The exposure time was 2.5 s, with frames collected for a total of 50 frames (0.05 s/frame) per sample. The dose rate was 21.2 e/pixel/s. SerialEM 3.7 was used for semiautomatic data acquisition.

### 4.4 Image processing and 3D reconstructions

The general strategy in the image processing follows the method in a hierarchical way as described.^49^ Data binned by 4 times is used for micrograph screening and particle picking. The data with 2-time binning is used for particle screening and classification. The particle after initial cleaning was subjected to extraction from the original clean micrograph and the resultant dataset was used for final cleaning and reconstruction. Raw movie frames were aligned with MotionCor2^50^ using a 9 × 7 patch and the contrast transfer function (CTF) parameters were estimated using Gctf and ctf in JSPR.^51^ Only the micrographs with consistent CTF values including defocus and astigmatism were kept for following image processing. For HCAR2-Gi1-scFv16 protein, 3,456 movies were processed by cryoSPARC v4.1.1.^52^ Each movie stack was aligned with patch motion correction. A total of 2,947,103 particles were extracted with auto-picking. After three rounds of 2D classification, the number of good particles was reduced to 515,376. The number of particles was further reduced to 311,666 by 3D classification and Ab-initio reconstruction. A 3.28 Å resolution density map at FSC 0.143 was obtained when the initial map of the particles was processed with homogeneous refinement, non-uniform refinement, and local refinement. For HCAR2-Gi1-scFv16 protein with niacin, 2,961 movies were processed by cryoSPARC v4.1.1. Each movie stack was aligned with patch motion correction. A total of 3,191,801 particles were extracted with the auto-picking. After three rounds of 2D classification, the number of good particles was reduced to 1,563,889. The number of particles was further reduced to 879,036 by 3D classification and Ab-initio reconstruction. A 2.69 Å resolution density map at FSC 0.143 was obtained when the initial map of the particles was processed with homogeneous refinement, non-uniform refinement and local refinement. For HCAR2-Gi1-scFv16 protein with acipimox, 1,706 movies were processed by cryoSPARC v4.1.1. Each movie stack was aligned with patch motion correction. A total number of 1,328,380 particles were extracted with the auto-picking. After three rounds of 2D classification, the number of good particles was reduced to 454,521. The number of particles was further reduced to 221,940 by 3D classification and Ab-initio reconstruction. A 3.23 Å resolution density map at FSC 0.143 was obtained when the initial map of the particles was processed with homogeneous refinement, non-uniform refinement and local refinement. For HCAR2-Gi1-scFv16 protein with MK-6892, 2,518 movies were processed by cryoSPARC v4.1.1. Each movie stack was aligned with patch motion correction. A total number of 1,988,362 particles were extracted with the auto-picking. After three rounds of 2D classification, the number of good particles was reduced to 558,862. The number of particles was further reduced to 291,441 by 3D classification and Ab-initio reconstruction. A 3.25 Å resolution density map at FSC 0.143 was obtained when the initial map of the particles was processed with homogeneous refinement, non-uniform refinement and local refinement.

### 4.5 Model building and refinement

Alphafold (https://alphafold.ebi.ac.uk/) was used to predict the human-HCAR2 structure, which was used as a template to build the HCAR2-Gi1-scFv16 complex model. Gi-scFV16 was built using the Gi1 heterotrimer from the FPR2-Gi cryo-EM structure (PDB 6OMM) as the template.^53^ All models were subsequently docked into the density maps using UCSF Chimera, followed by iterative manual adjustment and rebuilding in COOT 0.9.7 and phenix.realspace refinement. The final refinement model statistics were validated by Phenix. Model docking was carried out using MOE2019.01 software. The binding free energy was determined by Prime-MMGBSA in Maestro. The molecular graphics figures were presented using UCSF Chimera, UCSF ChimeraX, and PyMOL. The final refinement statistics were validated using Molprobity and shown in Supplementary information, Table S1.

### 4.6 cAMP assay

In order to measure the G-protein activation level, the cAMP-Gi kit (Perkin Elmer, TRF0263) was used. Wild-type HCAR2 and its mutants were cloned into a pcDNA3.1 vector. Before transfection, HEK-293 cells (ATCC CRL-1573) were seeded in 24-well culture plates at a density of 70-90% cells per well. Then the cells were transiently transfected with the plasmid using Lipofectamine 3000 reagents (Invitrogen, L3000). After 36 h, the culture media was removed, and the cells were washed with PBS buffer. The transfected cells were then plated into 384-well plates (4000/well) in a stimulation buffer and treated with 20 μM forskolin, 500 μM IBMX, and the test agonist for 30 min at 37°C. Thereafter, 5 μl cAMP Eu-cryptate reagent and 5 μl of anti-cAMP-d2 working solution were added to the 384-well plates and incubated for 1 h.^54^ Fluorescence signals were detected at 620/665 nm using the Multimode Plate Reader (PerkinElmer EnVision 2105).^55^ Data were analyzed with GraphPad Prism 9.0. The experiments were performed in triplicate.

### 4.7 Cell surface expression testing

Expression levels of HCAR2 plasmid in HEK-293 cells were determined by flow cytometry analysis and used to normalize the cAMP level. Specifically, the transfected cells were blocked with 5% BSA at room temperature for 15 min and labeled with anti-FLAG antibody (1:100, Thermo Fisher) at 4°C for 1 h. After washing with PBS buffer, the cells were incubated with anti-mouse Alexa 488-conjugated secondary antibody (1:300, Beyotime) at 4°C in the dark for 1 h. Each sample was counted with approximately 10,000 cellular events. The fluorescent intensity was quantified by a BD Accuri™ C6 Plus flow cytometer. The experiments were performed in triplicate. Values are presented as the mean ± SEM. Data were analyzed using GraphPad Prism 9.0, and significance accepted at *p* < 0.05.

## Acknowledgements

We would like to thank the Kobilka Cryo-Electron Microscopy Center, the Chinese University of Hong Kong, Shenzhen for our cryo-electron microscopy. This work was supported by funds from the Ganghong Young Scholar Development Fund, and the Shenzhen-Hong Kong Cooperation Zone for Technology and Innovation (HZQB-KCZYB-2020056). Y.D. is supported by grants from National Natural Science Foundation (General Project 32271263), Science, Technology and Innovation Commission of Shenzhen Municipality (Project code JCYJ20200109150019113; JCYG 20220818103009018), and in part by the Kobilka Institute of Innovative Drug Discovery at the Chinese University of Hong Kong, Shenzhen. X.P. is supported by grant from the National Natural Science Foundation of China (82000257).

## Author Contributions

Y.D., and J.X. conceived the study, and supervised the work; X.P., F.Y., P.N., and Z.Z. performed the experiments.; B. Z., G.C., W.G., C.Q., Z.W., and J.L. assisted in some experiments; X. P. and F.Y. prepared the manuscript; All authors discussed and analyzed the data; Y.D., X.P., F.Y., K.G., and J.X. revised the manuscript.

## Data Availability Statement

The data supporting this study are available from the corresponding author upon reasonable request. The 3D cryo-EM density maps of the niacin-, acipimox-, MK-6892-bound and apo HCAR2-Gi-scFV16 complexes have been deposited in the Electron Microscopy Data Bank database under accession codes EMD-35483, EMD-35484, EMD-35485 and EMD-35463, respectively. The atomic coordinates for the atomic models of the niacin-, acipimox-, MK-6892-bound and apo HCAR2-Gi-scFV16 complexes generated in this study have been deposited in the Protein Data Bank database under accession codes 8lJA, 8lJB, 8IJD and 8lJ3, respectively.

